# Enhanced EEG Forecasting: A Probabilistic Deep Learning Approach

**DOI:** 10.1101/2024.01.16.575836

**Authors:** Hanna Pankka, Jaakko Lehtinen, Risto J. Ilmoniemi, Timo Roine

## Abstract

Forecasting electroencephalography (EEG) signals, i.e., estimating future values of the time series based on the past ones, is essential in many real-time EEG-based applications, such as brain–computer interfaces and closed-loop brain stimulation. As these applications are becoming more and more common, the importance of a good prediction model has increased. Previously, the autoregressive model (AR) has been employed for this task — however, its prediction accuracy tends to fade quickly as multiple steps are predicted. We aim to improve on this by applying probabilistic deep learning to make robust longer-range forecasts.

For this, we applied the probabilistic deep neural network model WaveNet to forecast resting-state EEG in theta- (4–7.5 Hz) and alpha-frequency (8–13 Hz) bands and compared it to the AR model.

WaveNet reliably predicted EEG signals in both theta and alpha frequencies over 100 ms ahead, with mean errors of 0.8±0.6 µV (theta) and 0.7±0.5 µV (alpha), and outperformed the AR model in estimating the signal amplitude and phase. Furthermore, we found that the probabilistic approach offers a way of forecasting even more accurately while effectively discarding uncertain predictions.

We demonstrate for the first time that probabilistic deep learning can be utilised to forecast resting-state EEG time series. In the future, the developed model can enhance the real-time estimation of brain states in brain–computer interfaces and brain stimulation protocols. It may also be useful for answering neuroscientific questions and for diagnostic purposes.

## 1 Introduction

Electroencephalography (EEG) is widely used for non-invasively monitoring the neuronal activity of the brain. An important subfield of EEG signal analysis is the forecasting task, where one or more future values {*x_t_, …, x_t_*_+*m*_} of the EEG time series are estimated based on the previous values {*x_t−n_, …, x_t−_*_1_} of the same series. This problem has been studied at least since the 1990s (Blinowska & Malinowski, 1991) and it has recently gained more attention because of the growing demand for real-time EEG. Real-time EEG is used in applications such as brain–computer interfaces (BCI) and closed-loop neurostimulation, where a prediction model is sometimes required to allow time for algorithmic decision-making. For example, brain stimulation can be guided with EEG by anticipating future brain states and timing the stimulation accordingly (see, for example, Gharabaghi et al., 2014; Stefanou et al., 2019; Zrenner et al., 2018). A good prediction model can, therefore, enhance the operation of such applications as well as provide valuable insights into the neuronal activity of the brain.

Previously, only a few methods for forecasting EEG have been explored. The predominant prediction method is a linear autoregressive model (AR), which was used already in the earliest studies (Blinowska & Malinowski, 1991) and to date, it has been a popular choice in practical applications. For example, Chen et al. (2013) and Zrenner et al. (2018) used it for phase-locked brain stimulation. A few other methods have also been proposed but in the context of mental imagery classification for BCIs. Examples include multilayer perceptrons (Coyle et al., 2005), adaptive neuro-fuzzy inference systems (ANFIS) (Hsu, 2010; Komijani et al., 2019), and Elman recurrent neural networks (Forney & Anderson, 2011), where a one-step-ahead prediction is produced for a further classification task. Additionally, a couple of forecasting studies have focused purely on determining the future phase of the signal but did not attempt predicting the signal amplitude (Mansouri et al., 2017; McIntosh & Sajda, 2020). Ergo, so far the AR model is the only method used for long-range forecasts of the whole signal, but its prediction error tends to grow fast as multiple steps are forecasted. This is not surprising, as EEG time evolution is difficult to forecast — the signals are high-dimensional and non-stationary and the relationship between the past and future values is not deterministic. Moreover, signals from other brain areas and external inputs add to the difficulty of predicting the signal in a particular EEG channel. This indicates that a more complex model, compared to a simple linear one, would be needed to achieve reliable long-range forecasts.

A prominent approach to forecasting EEG is deep learning. During the last two decades, deep-learning techniques have demonstrated their power in modelling various high-dimensional data sets. The success of these methods lies in their ability to extract meaningful features from multidimensional data, which is why they hold promise for modelling EEG signals as well. Indeed, many studies on deep learning for EEG data have already been published, although these approaches almost solely consist of different classification tasks (Roy et al., 2019). Nevertheless, the results have been promising and there’s also one study already applying deep learning for the forecasting task (Bhowmick et al., 2020), but, again, only predicting one time step forward. While these studies preliminarily support the applicability of deep learning for EEG forecasting, there is still a need to explore its capability in long-range predictions.

Additionally, we hypothesized that a probabilistic approach would be a better fit for EEG forecasting compared to the previous point forecasting methods. Currently, many state-of-the-art time series forecasting methods are probabilistic (see for example Rangapuram et al., 2018; Salinas et al., 2020), as they offer some desirable properties. Probabilistic models predict a distribution over the future value, rather than a single continuous value. This allows the creation of multiple predictions per input by sampling from the distribution, and furthermore, it can give an uncertainty estimation for the predictions. An uncertainty estimation is particularly important when decisions (e.g., when to stimulate the brain) are made based on the predictions, as we can then refrain from action if the model is unsure.

Therefore, we propose forecasting EEG time series with the probabilistic deep-learning model WaveNet (van den Oord et al., 2016). WaveNet was originally developed for generating raw audio signals and has demonstrated excellent performance in this task. We hypothesized that its success with audio signals could make it well-suited for predicting EEG signals as well.

In this paper, we employ the deep learning model WaveNet to forecast EEG time series in theta- (4–7.5 Hz) and alpha-frequency (8–13 Hz) bands as well as deterministic synthetic data and test its capacity to predict signal amplitude and phase as well as oscillation peaks. We also compare the results to the AR model as it has so far been the only method for forecasting EEG time series multiple time steps ahead (see for example Blinowska & Malinowski, 1991; Chen et al., 2013; Zrenner et al., 2018). In addition, we examine the differences in prediction accuracy within subjects and channels, and whether there is a difference between training the model with data from all channels compared to training with data from a single channel. Finally, we explore a way of utilizing WaveNet’s probabilistic nature for finding unreliable predictions.

We find that WaveNet is suitable for forecasting both theta- and alpha-frequency bands and that it outperforms the AR model in this. Additionally, we show that WaveNet’s probabilistic forecasts can be used to narrow down a sharper prediction and detect time points that are hard to predict.

## 2 Materials and methods

### 2.1 WaveNet

In 2016, van den Oord et al. (2016) introduced WaveNet, a fully probabilistic generative deep learning model for creating raw audio signals. WaveNet models time series with conditional probabilities, where the probability of each value *x_t_*in the series is conditioned on *n* previous values: *p*(*x_t_|x_t−n_, …, x_t−_*_1_). It models these relationships between past and future values with causal convolutional layers, where the causal property ensures that no future values are considered when predicting the next time step. The model architecture (see Figure 1a) consists of a stack of WaveNet blocks, where the output from the previous block is fed to the next one as well as to the rest of the network to produce the final output. The output is a set of categorical probability distributions that has the same time dimensionality as the input, meaning that for each input time series (*x_t−n_, …, x_t−_*_1_) we obtain the probability distributions for the next time steps (*x_t−n_*_+1_*, …, x_t_*) (see Figure 1b). The distributions are created with a softmax activation as the last layer in the network, and the model parameters are optimized to maximize the log-likelihood of the predictions.

**Figure 1:**
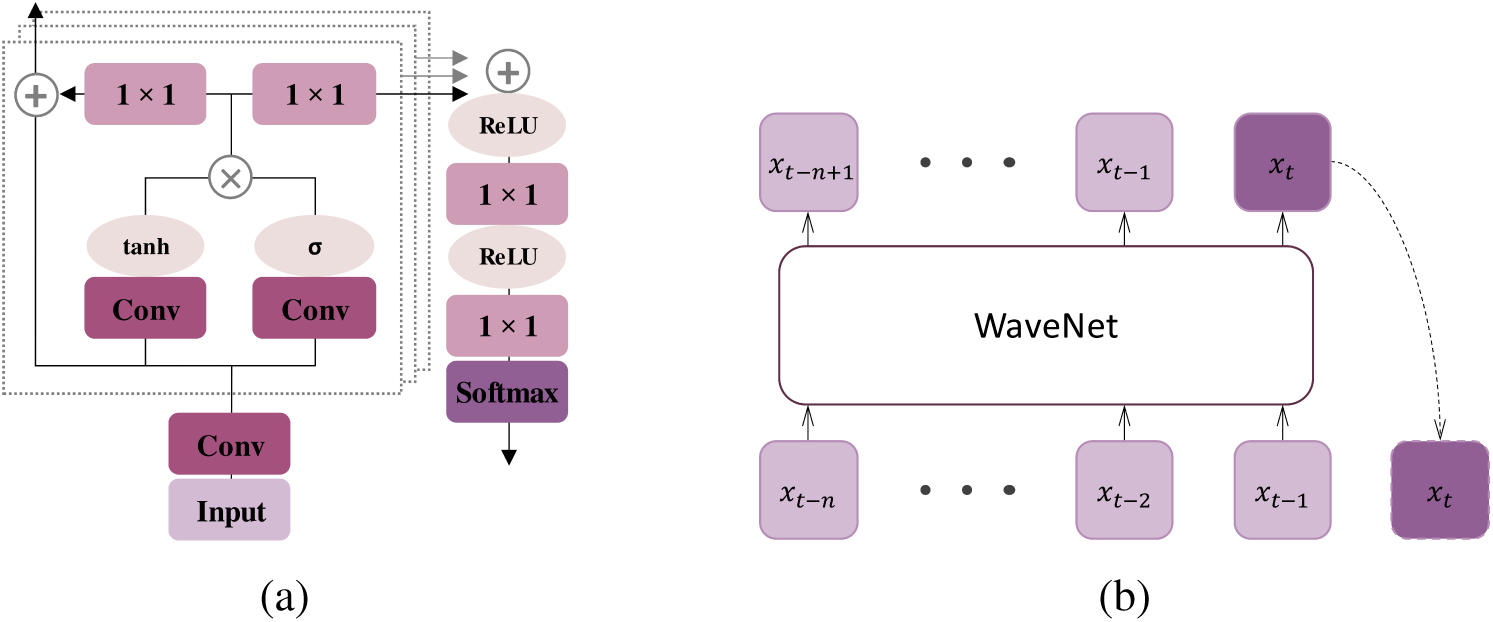
(a) Model architecture of WaveNet. WaveNet consists of two types of convolutional neural network layers: causal convolutions (“Conv”) and 1 *×* 1 convolutions, three types of activation functions: tanh, *σ*, and rectified linear units (ReLU) and a softmax layer. (b) WaveNet outputs a conditional probability distribution over the next value for each value in the input series. During training, the predictions can be generated in parallel and the loss can be calculated from the whole output sequence. At inference, predictions are generated one at a time and fed back to the network.

Modelling time series as categorical probabilities with convolution operations includes two challenges that WaveNet solves efficiently. First, time series like audio and EEG signals contain different structures at different time scales, which means that the model has to capture both small- and large-scale variations in the long temporal sequences. Achieving this with convolution operations can be computationally heavy because the receptive field has to be wide enough to cover the full range of these variations. To overcome this problem, WaveNet dilates the receptive field of the convolution operation by reading only every *d*th value. Throughout the model, this dilation factor *d* is exponentially increased in every WaveNet block as a power of two up until 512, after which it is repeated. This way we can achieve a receptive field size of 1024 with just 10 WaveNet blocks and a filter width of 2. The second challenge is that the resolution of digitized EEG signals typically ranges from 16 to 24 bits (Halford et al., 2016), which means that there are from 65,536 to 16,777,216 possible values for each time step. As computing probabilities for each of these values would not be feasible, the signal is quantized to 8-bit resolution to reduce the number of possibilities to 256. For a more detailed description of the model, we refer to the original article of van den Oord et al. (2016).

In this work, we adapted a version of the WaveNet model to be used for EEG time series forecasting using Tensorflow deep learning library (v 2.6.0) (Abadi et al., 2016).

### 2.2 Autoregressive model

The AR model of order *z* estimates the next value of a signal as the weighted sum of *z* previous values as follows:

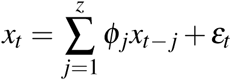

where {*x_t−_*_1_*, …, x_t−z_*} are the past *z* values, and {*φ*_1_*, …, φ_z_*} represent the coefficients of these past values. The coefficients can be determined in multiple ways; here, we apply the Yule–Walker equations (Walker, 1931; Yule, 1927), which estimate them with the auto-covariance of the time series. The coefficients are calculated from previous *z* time steps separately for each prediction interval (as opposed to fitting them for some data segment beforehand). A prediction of desired length is then created autoregressively, i.e., by adding the newest prediction to the sequence and using that sequence to generate the next one.

### 2.3 Synthetic data

We generated synthetic data for validating WaveNet in a fully deterministic scenario. Each sample was created as a product of two sine waves of different frequencies and with a random phase delay. The frequencies of the two waves were drawn at random from distinct normal distributions (with mean *µ* = 10 and standard deviation *σ* = 1 Hz and *µ* = 0.5*, σ* = 1 Hz), and the oscillation phase at the beginning of each wave was sampled from a uniform distribution [0°, 360°].

### 2.4 EEG data

We used resting-state EEG from 68 healthy subjects for training, validating and testing the model. The data were retrieved from an open online repository PRED+CT (http://predict.cs.unm.edu/, access number d003) (Cavanagh et al., 2019). There were recordings from 75 healthy subjects in total, but we ignored 7 of them due to missing or distorted segments. The measurements had been performed with a 60-channel Neuroscan system at the sampling rate of 500 Hz and with the reference channel between Cz and CPz channels. Each recording contained both eyes-open and eyes-closed conditions, 3 minutes each — for consistency, we chose to include only the eyes-open condition.

We tested WaveNet’s capacity on minimally processed data to keep the task as general as possible by not assuming any preprocessing pipelines or specific use cases. Therefore, no preprocessing apart from causal band-pass filtering was done. We band-pass filtered the EEG data to theta (4–7.5 Hz) and alpha (8–13 Hz) frequencies. To ensure that each time step in the filtered signal is independent of future time points, we chose to use a causal minimum-phase finite impulse response filter (order 825).

Before feeding the data to WaveNet, we quantized the signals to 8 bits. To ensure that this reduced signal resolution is sufficient for accurately representing the whole signal, we chose to present the model only with such signals, where the amplitudes were within the range of [*−*10, 10] µV. Following the original work of van den Oord et al. (2016), we also applied µ-law companding,

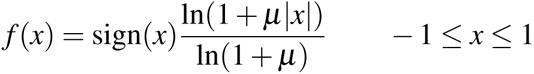

where *µ* = 255, before quantizing to introduce improved reconstruction compared to linear quantization. When making predictions with the AR model, we did not perform this quantization step.

### 2.5 Experimental conditions

We evaluated the performance of WaveNet under five experimental conditions. In the first condition, we trained WaveNet on synthetic data. The four other conditions were formed as combinations of two frequency bands and two data compositions. More specifically, WaveNet was trained on EEG time series that were a) band-pass filtered to either theta- (4–7.5 Hz) or alpha-frequency (8–13 Hz) band and b) selected from either all channels (All condition) or just the C3 channel (C3 condition). We tested the inclusion of data from all channels because this results in a larger and more diverse training set compared to a set from a single channel. This might be beneficial as the increased variety in the training set can prevent overfitting and help WaveNet extract features of the underlying data distribution. Notably, the model was not conditioned on the channel identity, meaning that it could not differentiate which channel each signal was coming from. On the other hand, we wanted to test whether WaveNet could more accurately predict a particular channel, had it seen data only from that channel. This might indeed be the case, as EEG signals originating from different parts of the brain tend to differ from each other — thus WaveNet might learn the features of a specific channel better if it was trained with data only from that channel. We chose to test this with the C3 channel because it has been previously used in the literature (see, for example, Zrenner et al., 2020).

### 2.6 Cross-validation

We performed 4-fold cross-validation to evaluate the performance of WaveNet on the whole dataset. Thus, the dataset of 68 subjects was divided into four parts, each containing data from 17 subjects. For each cross-validation iteration, one part was put aside for testing, while the rest of the subjects were used for the training process. Therefore, the training process included 51 subjects divided at random into training (47 subjects) and validation sets (4 subjects). This process ensures that neither the subjects in the training nor the validation sets were used in the testing phase for any of the cross-validation iterations.

This procedure was applied separately for each of the four experimental conditions where EEG data were used.

### 2.7 WaveNet’s hyperparameters

Training a deep-learning model requires selecting a set of hyperparameters — we chose some of these by hand and performed a grid search to optimise the rest. Our goal was not only to select hyperparameters that produce the most accurate predictions but also to choose them in a way that ensures the usability of the model in real-time applications.

The kernel width of the convolution operations and the number of WaveNet blocks were chosen by hand such that the resulting receptive field size is as small as possible whilst still covering enough signal to make reliable forecasts. Short inputs are preferred in many real-time applications, where long resting-state sequences might not be available — this might happen, for example, when there are stimuli disturbing the signal. In addition, wider receptive fields would prolong prediction times, which is not a desirable property for online predicting. Thus, we set the kernel width to two and the number of WaveNet blocks to 10. This combination produces a receptive field size of 1024, which we found to be sufficient for determining the next time step.

Next, we applied the grid search to find the optimal learning rate and the number of convolutional filters. We tested eight configurations, consisting of two learning-rate values ({0.001, 0.0001}) and four convolutional filter counts: {16, 24, 32, 40} for the All condition and {8, 16, 24, 32} for the C3 condition. We chose the parameters that produced the lowest validation loss after 600 epochs. Notably, the grid search was performed separately for each cross-validation group to avoid leaking properties of the test data to the training process.

For training WaveNet with synthetic data, we used the same hyperparameters that were chosen for models in the alpha band in the All condition.

### 2.8 Training WaveNet

We trained WaveNet with samples of either synthetic data or EEG time series. The division of data into training, validation, and test sets is described in Section 2.6. We chose 2250 training and validation examples of length 2000 at random from each subject in the training and validation sets. When training models with the All condition, each sample was chosen at random from the 60 channels, whereas for the C3 condition, only the C3 channel was used. From those examples, we then discarded samples that contained amplitude values outside our target range [*−*10, 10] µV.

We then trained WaveNet for 600 epochs with a batch size of 32. We chose new training examples at the beginning of each epoch to avoid overfitting the model by introducing variation to the samples. We used an Adam optimizer (Kingma & Ba, 2014) with a decaying learning rate to minimize the categorical cross-entropy loss between predicted and target labels. For calculating the loss, we only included time points whose receptive field did not contain any padding zeros.

### 2.9 Order of the autoregressive model

We optimized the AR model’s order with the same validation data sets that were used to select WaveNet’s hyperparameters. To allow full comparability with the WaveNet model, we used the same number of time steps to calculate the model coefficients as was selected as the receptive field for WaveNet. Thus, we selected 300 random samples of length 1024 from each of the 4 subjects in each validation set and excluded the samples containing values outside the range [*−*10, 10] µV. For each of the samples, we created predictions of length 75 (150 ms) with 30 distinct model orders: {10, 20*,…,* 300}. We then calculated the mean average error over the whole prediction interval. Finally, for each cross-validation group, we chose the model order that produced the lowest mean absolute error over the whole validation set of the group.

### 2.10 Performance evaluation

We evaluated WaveNet’s prediction performance with synthetic data and with EEG time series from the 17 subjects left out of the training set of each cross-validation group. To evaluate WaveNet’s performance under the All condition and to compare that to the AR model, we composed a test set by selecting 120 samples at random from every channel of each subject. Another test set was created to compare the All and C3 conditions: here, we selected 1500 random samples from the C3 channel of each subject. For both of these test sets, as well as for evaluation with synthetic data, we created forecasts of length 75 (150 ms). Additionally, we determined how well WaveNet under All condition can detect high and low peaks occurring 50 ms in the future compared to the AR model. For this peak detection task, we made a third test set, for which we chose inputs from all of the channels such that 50 ms forwards from each sample there was a high or low peak in the measured signal. To detect the peaks, we generated forecasts of length 50 (100 ms), meaning that the peak was expected to appear in the middle of the predicted window. From all of the test sets, we dismissed the samples that were not in the target range [*−*10, 10] µV.

We generated 10 sampled predictions per input in the test sets — each sample was formed separately by predicting one time step at a time and feeding the result back into the network. At each time step, the predicted value was sampled with a temperature of 0.5 (Guo et al., 2017) by squaring the probabilities in the output distribution. This stretching of the output distribution was done, as we found in our initial testing that it produced better predictions compared to sampling from the original output distribution. In all of the comparison experiments, the final forecast was calculated as the mean of the 10 sampled predictions. Additionally, we tested whether WaveNet’s prediction performance could be improved by factoring in the uncertainty in the predictions. We hypothesized that, rather than averaging all the samples, the forecasts could be made sharper by including only the samples that agree enough with each other. To test this, we used the same 10 samples as before but applied the following rules at each predicted time step: Find the largest group with at least *s* samples that are at most *r* µV apart from each other and form a prediction as a mean of the samples in this group. If there is no such group, refrain from predicting. We tested this with group sizes {4, 5, 6} and ranges {0.5, 0.75, 1} µV.

For some of the analyses, we wanted to determine the phase of the signal. This was performed by calculating the analytic signal using the Hilbert transform and then extracting the angles of the resulting complex numbers. Due to the edge effect caused by the Hilbert transform, we discarded the last 10 ms of each phase estimation.

Lastly, we performed statistical testing to determine the statistical significance of the differences we found. For comparing group means, we used a *t*-test — when Student’s *t*-test’s assumption of equal variances was not met, we used Welch’s *t*-test. For comparing the differences in variances, we applied Levene’s test.

## 3 Results

### 3.1 Grid search

In each cross-validation group, the best-performing model had a learning rate of 0.001. There was more variation in the number of convolution filters in the best-performing models, although within each experimental condition the variation was small. In the All condition, 40 filters produced the lowest validation error in each cross-validation group apart from one group in the theta band, where the model with 32 filters performed the best. In the C3 condition, the grid search resulted in 24 convolution filters for one of the cross-validation groups in the theta band and for three groups in the alpha band. Conversely, 16 filters were chosen for three groups in the theta and one group in the alpha band.

### 3.2 Orders of the autoregressive model

The chosen model orders were similar for each cross-validation group (see Figure 2). The lowest mean absolute error during the 150-ms prediction interval was achieved with model order 100 in three cross-validation groups in the theta band and one in the alpha band. Furthermore, for the fourth group in the theta band, model order 120 and for the remaining three groups in the alpha band, orders 110, 120 and 170 were chosen.

**Figure 2:**
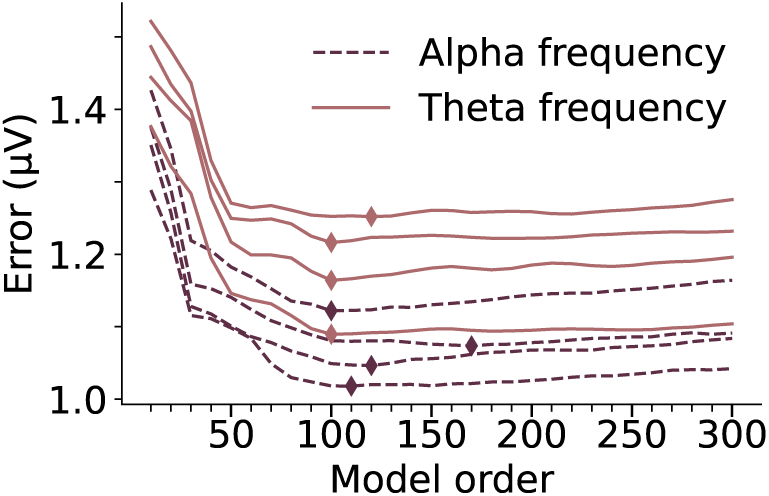
Results of optimization of the AR model order. The lines show the mean absolute error during a 150-ms prediction interval in each cross-validation group in theta- (violet) and alpha-frequency (reddish brown) bands. The chosen model orders with the lowest error are marked with a diamond shape.

### 3.3 Synthetic data

WaveNet reliably learned the features of the synthetic data. Figure 3 shows representative examples of forecasts made with test samples. The 10 predicted samples have minimal variance amongst each other and they closely follow the original signal. Thus, the representative power of the WaveNet model is sufficient for capturing the features of the deterministic data.

**Figure 3:**
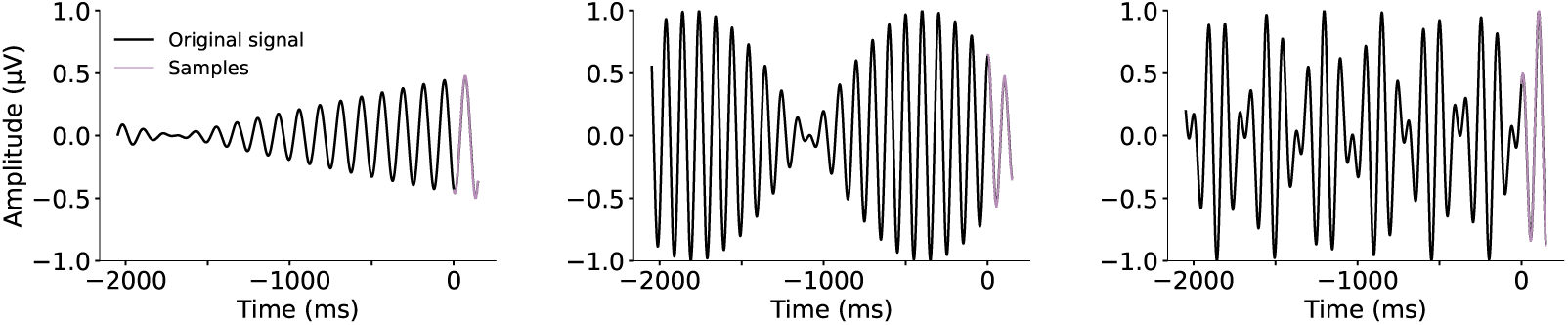
Example forecasts of length 150 ms with fully deterministic synthetic data. Each prediction interval contains 10 samples. As the predicted samples (in purple) are very close to the original signal, the original signal (in black) is not visible underneath the predicted signal.

### 3.4 Comparison of frequency bands

The prediction performances for the theta and alpha bands differed only slightly (Table 1) — Figure 4 shows the median prediction errors as a function of time while Figure 5 presents example predictions from both frequency bands. In the beginning, the prediction error for alpha frequencies increased faster than for theta, probably due to theta oscillations being slower — within the same amount of time, there is less change that needs to be predicted. Additionally, some of the differences in the error time series appeared to be due to the periodicity of the signals, which caused fluctuation according to the respective frequencies in both of the frequency bands.

**Figure 4:**
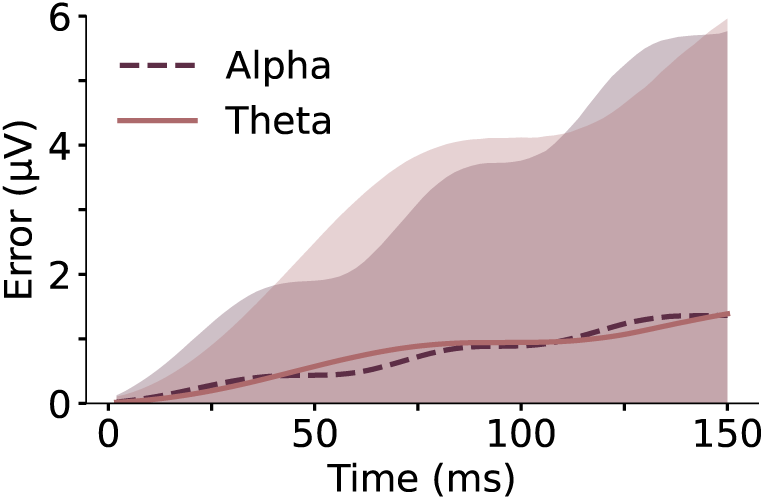
Median prediction error over time (dark lines) for theta- (4–7.5 Hz) and alpha-frequency (8–13 Hz) bands. The shaded area (lighter colour for the theta and darker for the alpha band) represents the range of these errors with the top 2.5% of outliers cut out.

**Figure 5:**
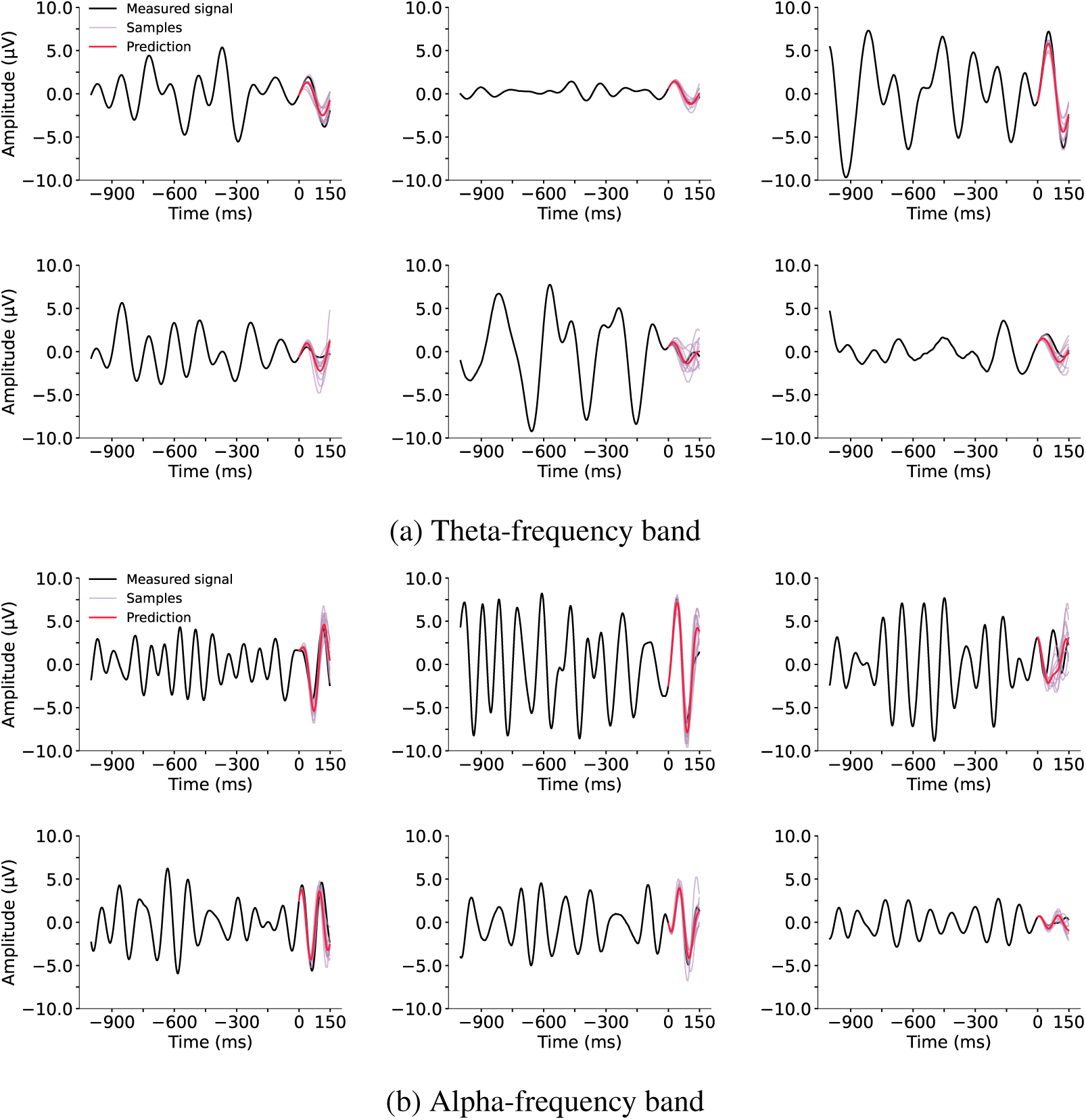
Example forecasts from the All condition, where the model was trained and tested with data from all of the 60 channels. Each prediction (in red) is formed as the mean of 10 samples (in purple). The measured signal is presented in black.

**Table 1:** Prediction errors (mean ± standard deviation) of WaveNet and the AR model. All of the differences in means were statistically significant (Welch’s t-test: *t*(539162) = *−*109.4 and *t*(532394) = *−*114.7 for amplitude errors and *t*(544430) = *−*84.9 and *t*(544639) = *−*73.0 for phase errors in theta and alpha frequency, respectively. *p < .*05 for all).

| | Amplitude ($\mu\text{V}$) | | Phase ($^\circ$) | |
| --- | --- | --- | --- | --- |
| | $\theta$ | $\alpha$ | $\theta$ | $\alpha$ |
| **WaveNet** | $0.8 \pm 0.6$ | $0.7 \pm 0.5$ | $24.7 \pm 23.3$ | $24.2 \pm 23.2$ |
| **AR** | $1.0 \pm 0.7$ | $0.9 \pm 0.6$ | $30.5 \pm 27.3$ | $29.0 \pm 26.2$ |

### 3.5 Comparison to the autoregressive model

Figure 6 shows the median amplitude errors of WaveNet and the AR model and the corresponding median phase errors. Table 1 summarizes the means of these prediction errors. For both frequency bands, the median prediction error during the whole prediction interval was consistently lower for WaveNet than for the AR model. In the theta band, the median prediction error of WaveNet did not exceed 1 µV until 114 ms, whereas the median error of the AR model exceeded 1 µV after 66 ms. Respectively, in the alpha frequency, the 1-µV error was exceeded at 108 ms with WaveNet and at 94 ms with the AR model. Thus, WaveNet’s prediction reliability remained high for considerably longer than that of the AR model.

**Figure 6:**
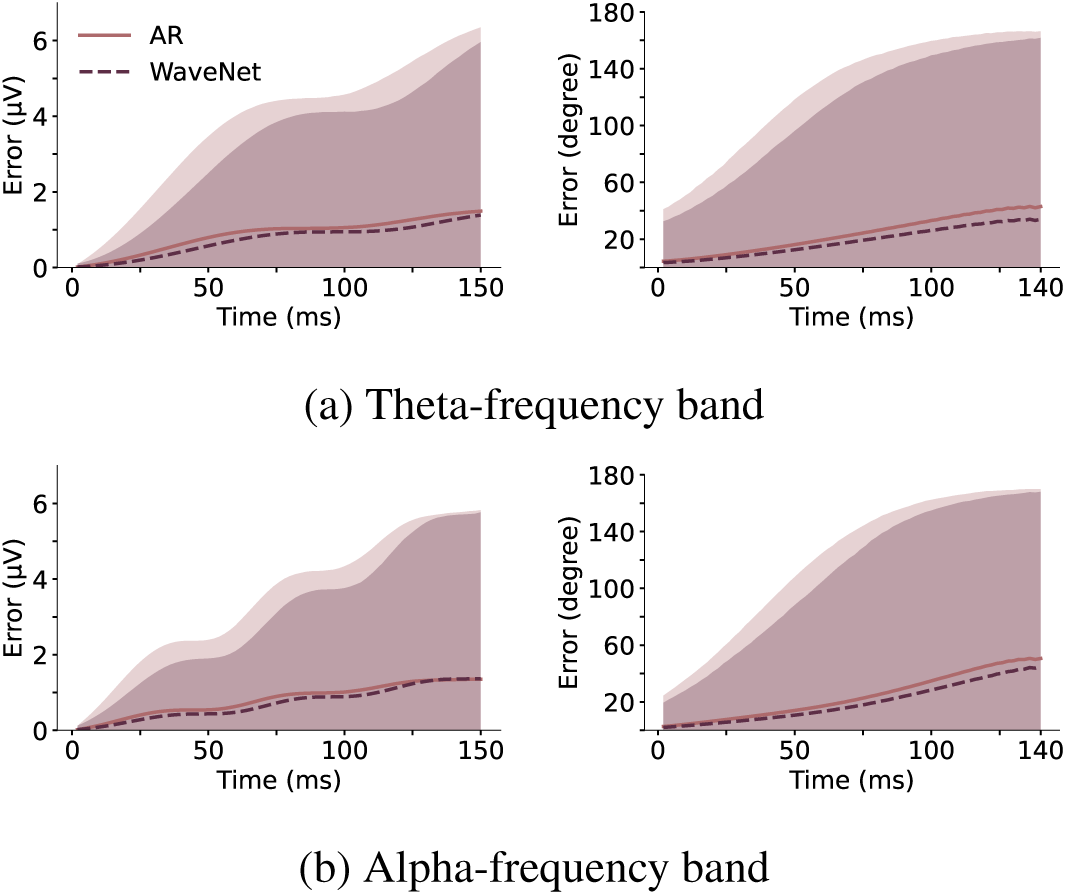
WaveNet’s forecasting performance compared to the autoregressive model. Median errors in amplitude (left) and phase are presented with dark lines. The coloured area represents the range of these errors with the top 2.5% of outliers cut out. The slight distortion at the end of the phase error sequences is caused by the edge effect of the Hilbert transform. WaveNet’s median prediction errors remained lower than those of the AR model during the whole prediction interval.

The results of the phase detection task are presented in Figure 7; its median errors, as well as the lower and upper quartiles, are displayed in Table 2. The AR model’s phase predictions were less precise than WaveNet’s predictions, of which 72% were within 20 degrees of the target phase, whereas only 65% of the AR model’s predictions were within that distance. What stands out from the distributions of the predicted phases is that in the alpha frequency, the trend lines of WaveNet’s and AR model’s predictions are behind the target phase, whereas in the theta frequency, they are ahead of the target phase. This means that in the alpha frequency, both models are slightly prone to predict the frequency to be too low, whereas, in the theta frequency, they are both more inclined to estimate the frequency to be too high rather than too low.

**Figure 7:**
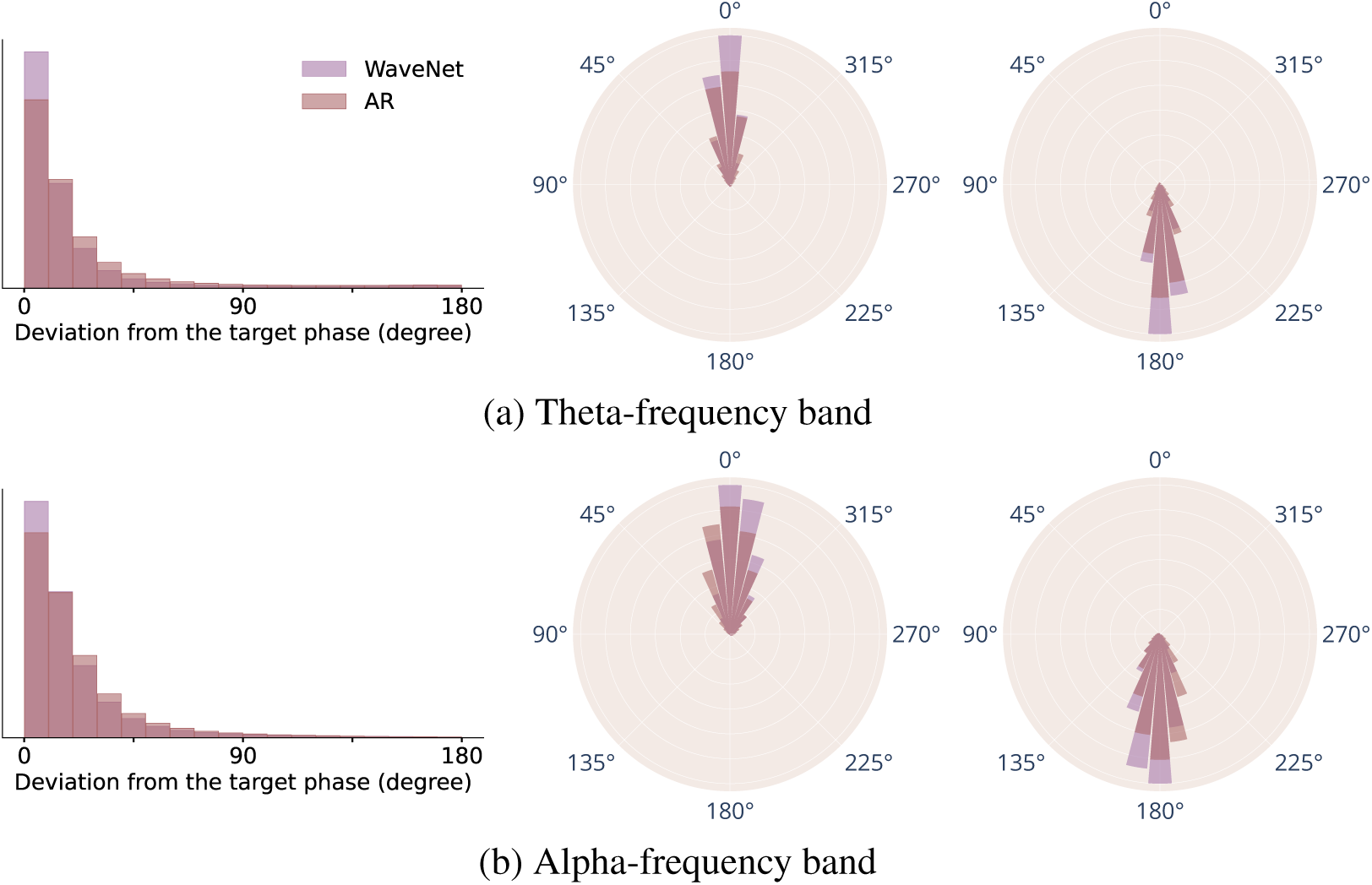
We found that WaveNet outperforms the AR model in estimating peaks that occur 50 ms in the future. The histograms on the left present how the deviations from the target phases were distributed between the models. The rose plots on the right display the predicted phases for the high (left) and the low peaks (right) with both models: the radial extent of each bar shows how many of the predictions were within the phase range indicated by the width of the bars.

**Table 2:**
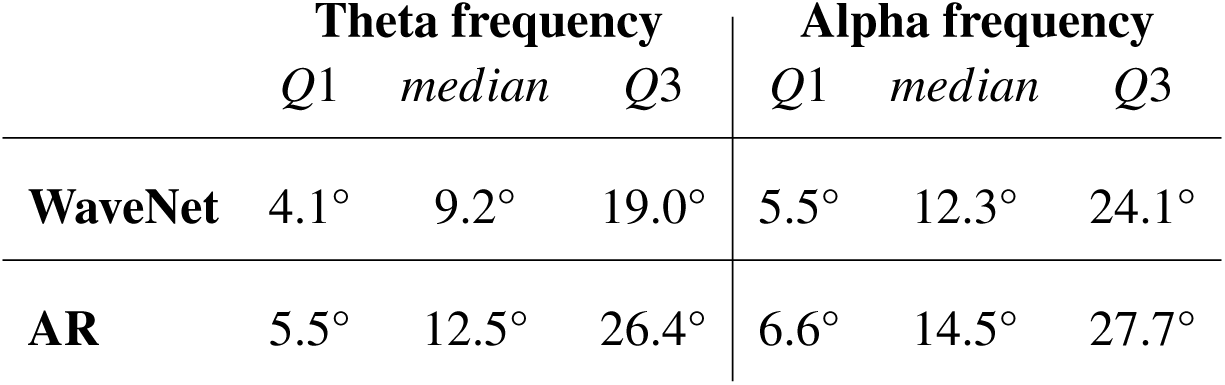
Deviation from the target phase in the peak-prediction task.

### 3.6 Differences between cross-validation groups, subjects and channels

We analysed whether WaveNet’s forecasting performance varied within cross-validation groups, individual subjects, and channels. We found the distributions of median prediction errors to be similar across all cross-validation groups for both frequency bands. This suggests that WaveNet has the capacity to robustly learn generalisable features from the different training data sets.

When comparing the median prediction errors of channels versus subjects, we found that there was more variation within channels than subjects when predicting the amplitude (Figure 8a). However, we found the opposite with the phase estimates of the same predictions (Figure 8b). This might be caused by the variation in amplitude sizes between channels — the closer the channels are located to the reference channel, the smaller their amplitudes are. For this reason, it is expected that the absolute errors in amplitudes are also smaller the closer we go to the reference channel. To further investigate this, we calculated how large the amplitude errors are relative to the measured signal (Figure 8c), observing that this, indeed, resulted in error distributions similar to those of the phase errors. Thus, the prediction results varied more between subjects than between channels when taking into account the variation in amplitude sizes between channels.

**Figure 8:**
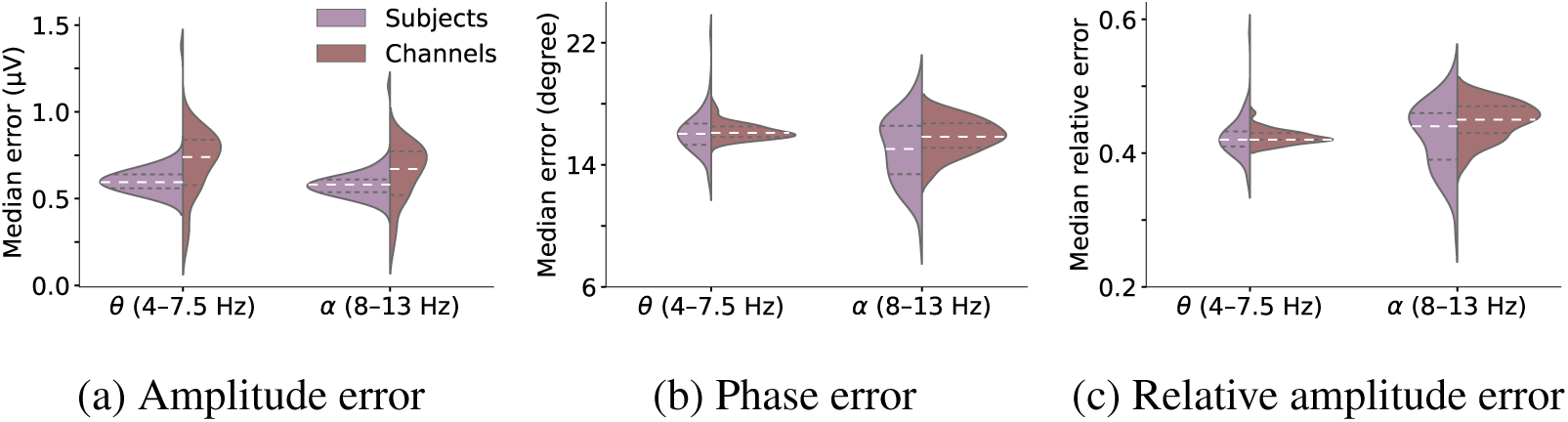
Error distributions of median errors of subjects versus channels: each data point is the median error of one subject or channel. The white dashed lines show the medians and the grey dashed lines the lower and upper quartiles of each distribution. All of the differences in group variances were statistically significant (Levene’s test: *F*(1, 126) equals 17.2 and 20.7 for amplitude, 16.3 and 30.9 for phase, and 17.1 and 16.1 for relative amplitude errors in theta and alpha frequencies, respectively. *p < .*05 for all).

### 3.7 Limiting training data to a single channel

We compared WaveNet’s ability to forecast the C3 channel under the C3 and All conditions. We calculated the median error of each 150 ms forecast and found that the means of these errors were nearly equal for both conditions: 0.72 *±* 0.48 µV (All condition) and 0.69 *±* 0.47 µV (C3 condition) in the theta-frequency band and 0.62 *±* 0.36 µV (All condition) and 0.61 *±* 0.36 µV (C3 condition) in the alpha-frequency band. The differences in means were statistically significant but of no practical significance (Welch’s *t*-test: *t*(178764) = 12.2*, p < .*05 in theta frequency and *t*(140404) = 5.1*, p < .*05 in alpha frequency).

### 3.8 Predicting only with samples that agree with each other

We experimented with utilizing the probabilistic nature of WaveNet to further improve its prediction accuracy. Figure 9a shows how the median error of the predictions is dependent on the minimum group size and maximum range of the generated samples. Almost all of the conditions produce more accurate predictions compared to the original predictions, where we always use all of the samples.

**Figure 9:**
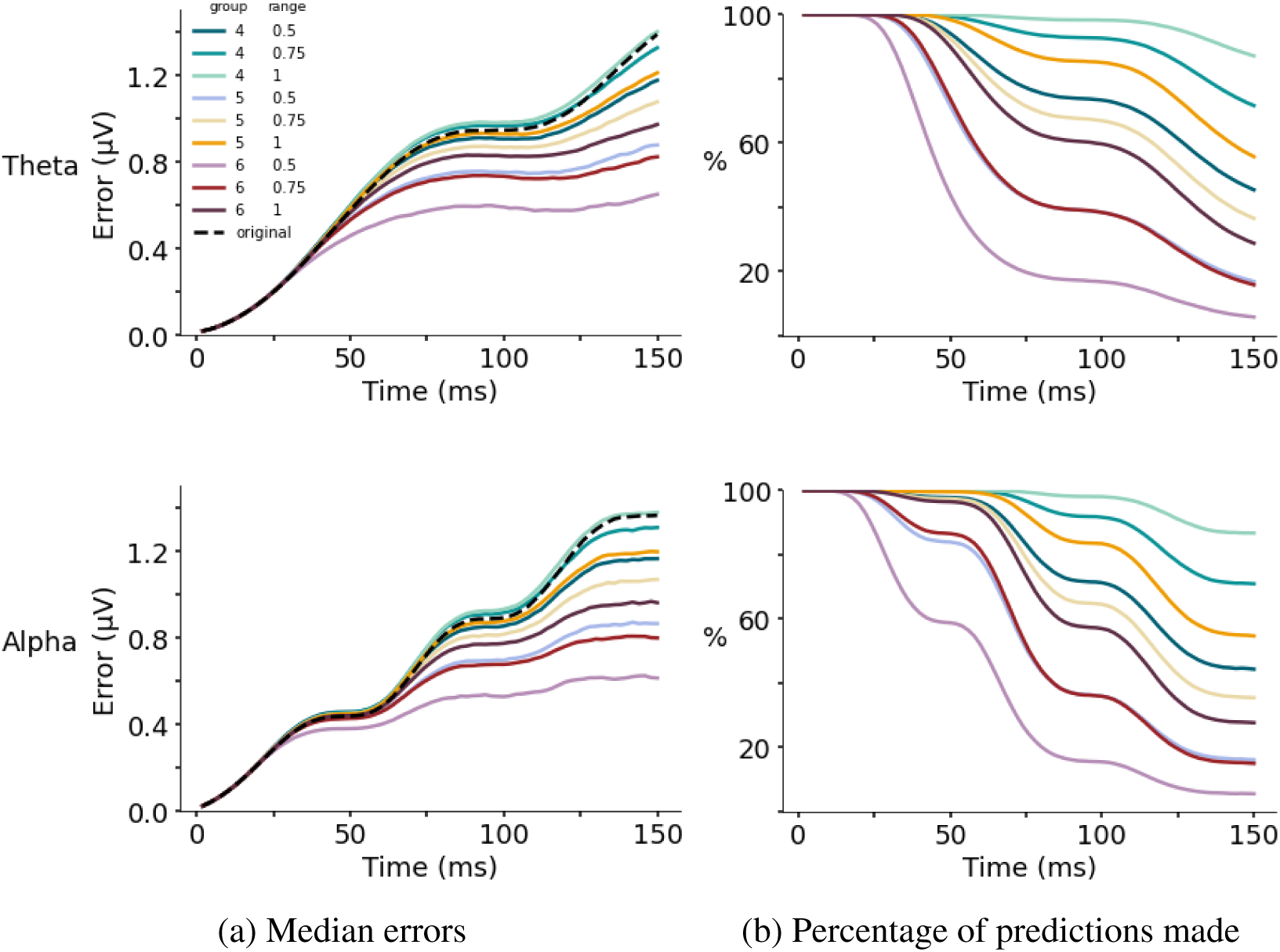
Results of using only samples that agree with each other and refraining from predicting, if such samples are not found. Each condition has a minimum group size and a maximum range. The former specifies the minimum number of samples that must be within a defined range at each time point to make a prediction, while the latter defines this range. For example, with *group* = 4 and *range* = 0.5, there has to be 4 or more samples within 0.5-µV range and the prediction is defined as the mean of all of the samples (*≥* 4) that fit into this range. If no such group is found, we refrain from predicting. The figures on the left show the median errors of each group size and range combination as functions of time in theta (upper) and alpha bands. The figures on the right show the percentage of predictions that were made at each time point.

The lowest median errors were achieved in both frequency bands with the strictest conditions: {6, 0.5}, {6, 0.75}, and {5, 0.5} for group size and range (µV), respectively. Correspondingly, fewer predictions were made with these strictest conditions compared to the other conditions (Figure 9b). This implies that excluding outlier samples yields sharper forecasts and that scattered samples can identify points that are difficult to predict.

## 4 Discussion

We investigated the use of probabilistic deep learning for forecasting multiple steps of resting-state EEG time series in theta-and alpha-frequency bands and found WaveNet model to be highly suitable for this task. WaveNet was able to forecast both frequency bands over 100 ms with less than 1-µV median absolute error. Moreover, it was able to predict deterministic simulated data visually perfectly. This indicates that a significant part of the prediction error with real data originates from its stochastic properties.

We found that WaveNet outperformed the linear AR model in forecasting the amplitude and phase of EEG signals as well as in detecting oscillation peaks. Notably, this was the case even though we optimized the AR model order, which led to a significantly improved prediction accuracy compared to a lower-order AR model reported in previous studies (Chen et al., 2013; Zrenner et al., 2018). Nevertheless, these results imply that a deep learning approach is, as hypothesized, more apt at modelling high-dimensional and non-stationary EEG signals compared to simpler linear models.

We also found that WaveNet’s prediction accuracy varied more between subjects than between channels when taking into account the variation in amplitude sizes between channels. These differences between subjects may be due to differences in brain function or differences between recordings. To investigate whether the differences are truly between subjects or, instead, between measurements, we would need to compare multiple recording sessions from a single subject. We could not test this as we only had one recording session per subject. This should therefore be further explored in future studies — if there is real variation in the input–output relationships between subjects, the prediction accuracy could be improved by conditioning WaveNet on subject identity. This approach was suggested also in the original WaveNet paper (van den Oord et al., 2016), where they found the model to perform better when trained on multiple speakers and conditioning on speaker identity compared to training on a single speaker.

Furthermore, we tested if limiting the training data to a single channel, C3, affects the forecasting performance but found the effect to be minute. Based on this result, there is no considerable benefit in training WaveNet with data from only one channel. The findings here further demonstrate that the differences between channels are not big; thus, limiting the channels that the model sees does not substantially improve the forecasting performance on one channel. However, the small improvement in accuracy suggests that conditioning WaveNet on channel information might improve performance; by conditioning, we get both the benefits of the diverse training set and the information about the characteristics of each channel.

Finally, we introduced one way of using WaveNet’s probabilistic predictions as an uncertainty measure: by forming the prediction by including only samples that are close enough to each other, we were able to decrease the prediction error. We found that the stricter the rules were, i.e., the more samples were required and the closer they needed to be, the sharper the forecasts were. However, this also went hand in hand with the amount of inputs we weren’t able to form any prediction for. Hence, with this approach, there emerges a tradeoff between the accuracy of the forecast estimation and how frequently we can predict. It is then dependent on the use case, how these factors should be weighted.

However, WaveNet also has drawbacks that could be addressed in future research. Firstly, WaveNet needs a considerably long input, which might be a disadvantage, especially with frequent stimuli present. Two main solutions could be to either try to shorten the input or examine whether WaveNet could make reliable forecasts regardless of a stimulus disturbing a part of the input. Secondly, WaveNet is drastically slower in creating predictions compared to the AR model. In real-time applications, a parallel WaveNet (van den Oord et al., 2018) could then be used; the parallel implementation was reported by the authors to be three orders of magnitude faster than the original. Lastly, we limited WaveNet to model band-pass filtered amplitudes in the range of [*−*10, 10] µV. In real-time forecasting, in case amplitudes outside this range are present within the current receptive field (i.e. past 1024 time steps), the model would not be able to predict and would need to wait until the signal is again within the range. Future research could look for ways to expand the range, in case there is a need to predict amplitudes outside this range. However, if values outside this range are specific to artefacts or otherwise low-quality data, it may be beneficial to refrain from predicting in these cases.

Future research could also look into modelling other frequency bands than theta and alpha. Additionally, WaveNet could be tested with, for example, patient data to discover potential group-specific differences in forecasting performance.

## 5 Conclusion

We investigated using the probabilistic deep learning model WaveNet for forecasting resting-state EEG time series multiple steps into the future. The novelty in this approach compared to previous methods is twofold: the deep learning architecture can capture the complex nature of the signals and the probabilistic approach provides an uncertainty measure for the forecasts.

Our results show that WaveNet is well-suited for forward predicting EEG and yields improved outcomes compared to the state-of-the-art linear AR model. Moreover, our findings suggest that conditioning WaveNet on subject or channel identity or both could further improve its performance. In the future, this model can improve real-time estimation of brain states in brain–computer interfaces and brain stimulation protocols. Moreover, the model may help answer novel neuroscientific questions and has potential applications in developing diagnostic and prognostic tools for various brain disorders.

## Acknowledgements

This project has received funding from the European Research Council (ERC Synergy) under the European Union’s Horizon 2020 research and innovation programme (ConnectToBrain; grant agreement No 810377). Hanna Pankka received support from the Jenny and Antti Wihuri Foundation. Jaakko Lehtinen was supported by the European Research Council (ERC Consolidator Grant 866435).

